# Across-subjects classification of stimulus modality from human MEG high frequency activity

**DOI:** 10.1101/202424

**Authors:** Britta U. Westner, Sarang S. Dalal, Simon Hanslmayr, Tobias Staudigl

## Abstract

Single-trial analyses have the potential to uncover meaningful brain dynamics that are obscured when averaging across trials. However, low signal-to-noise ratio (SNR) can impede the use of single-trial analyses and decoding methods. In this study, we investigate the applicability of a single-trial approach to decode stimulus modality from magnetoencephalography (MEG) high frequency activity. In order to classify the auditory versus visual presentation of words, we combine beamformer source reconstruction with the random forest classification method. To enable group level inference, the classification is embedded in an across-subjects framework.

We show that single-trial gamma SNR allows for good classification performance (accuracy across subjects: 66.44 %). This implies that the characteristics of high frequency activity have a high consistency across trials and subjects. The random forest classifier assigned informational value to activity in both auditory and visual cortex with high spatial specificity. Across time, gamma power was most informative during stimulus presentation. Among all frequency bands, the 75-95 Hz band was the most informative frequency band in visual as well as in auditory areas. Especially in visual areas, a broad range of gamma frequencies (55-125 Hz) contributed to the successful classification.

Thus, we demonstrate the feasibility of single-trial approaches for decoding the stimulus modality across subjects from high frequency activity and describe the discriminative gamma activity in time, frequency, and space.

**Author Summary:** Averaging brain activity across trials is a powerful way to increase signal-to-noise ratio in MEG data. This approach, however, potentially obscures meaningful brain dynamics that unfold on the single-trial level. Single-trial analyses have been successfully applied to time domain or low frequency oscillatory activity; its application to MEG high frequency activity is hindered by the low amplitude of these signals. In the present study, we show that stimulus modality (visual versus auditory presentation of words) can successfully be decoded from single-trial MEG high frequency activity by combining source reconstruction with a random forest classification algorithm. This approach reveals patterns of activity above 75 Hz in both visual and auditory cortex, highlighting the importance of high frequency activity for the processing of domain-specific stimuli. Thereby, our results extend prior findings by revealing high-frequency activity in auditory cortex related to auditory word stimuli in MEG data. The adopted across-subjects framework furthermore suggests a high inter-individual consistency in the high frequency activity patterns.

## Introduction

Since the first reports of cortical gamma band activity [1, 2], these high frequency responses have been linked to a plethora of brain processes and mental tasks, for example visual perception and processing [3, 4, 5, 6], auditory perception [7, 8] or memory [9, 10, 11, 12]. Although numerous theories about the origin and function of these high frequency oscillations and their relation with lower frequencies like theta and alpha have been proposed (e.g., [13, 14, 15]), there is an ongoing debate about whether gamma band responses reflect narrowband oscillations or broadband power increases, possibly echoing an increase in spiking activity [16, 17, 6]. One obstacle in this quest is the 1/f characteristic of the brain’s frequency power spectrum and a low signal-to-noise ratio (SNR) of gamma band activity in MEG or electroencephalography (EEG) recordings. To increase SNR, trial averaging is a frequently used tool to cancel out random variance. However, this approach can potentially obscure or cancel meaningful brain activity [18]. Indeed, local field potentials and electrocorticographic data from monkeys revealed systematic trial-to-trial variations in gamma power and frequency in a visual [19] and a memory task [20]. Importantly, the averages across trials in these studies displayed the classic sustained gamma effect, indicating that single-trial responses are crucial to understand the brain’s dynamics [18]. One powerful approach to assess single-trial information are multivariate decoding techniques. Whether such methods are applicable on low SNR gamma band MEG data, however, remains unclear. In the present paper, we investigate the predictive value of single-trial gamma power regarding the modality of stimulus presentation (auditory or visual presentation of words) in human MEG data. While comparable contrasts have been used to test classifier performance or as example datasets (e.g., [21, 22]), our aim was to unravel single-trial high frequency patterns in human MEG data. To decode information about stimulus-modality from the time-frequency data, we used a combination of beamforming [23] and random forest classification [24]. This approach was embedded into an across-subjects cross-validation framework, where the classifier was tested on unseen subjects to assess the generality of the spatial time-frequency pattern. Our results confirm that gamma SNR in single trials is high enough to achieve stable classification accuracy significantly above chance. Interestingly, the classification model yields high informational value to a broad bandwidth in the gamma range. Furthermore, we show that the characteristics of the gamma activity are similar enough across trials and even subjects to yield reliable classification performance.

## Results

To assess the predictive value of single-trial gamma power towards stimulus modality, we used MEG data from 20 subjects and adopted an across-subjects classification scheme. Data was first source reconstructed with an linearly constrained minimum variance (LCMV) beamformer, subsequently, we used the random forest algorithm to classify the modality of stimulus presentation (auditory or visual).

The random forest model classified auditory versus visual trials with 66.44 % accuracy, which is significantly better than chance (binomial test, *n*_*trials*_ = 4270, *p* < 0.001). As the confusion matrix in Figure 1A shows, the accuracy was slightly better for auditory trials (69.60 %) than for visual trials (63.19 %). In the adopted 20-fold cross-validation scheme, every fold corresponded to the data of one subject, hence, the classifier was always tested on data of one subject which was not included in building the model. The classifier accuracy on the 20 cross-validation folds is depicted in Figure 1B. The performance on the different folds is diverse, ranging from 50.98 to 84.86 %, however, the accuracy for all but three folds is above 60 % (note that the folds, since they are part of the whole classifier model, are not tested for significance). The good classifier performance indicates that the gamma power patterns are remarkably stable across trials and even subjects.

**Figure 1:**
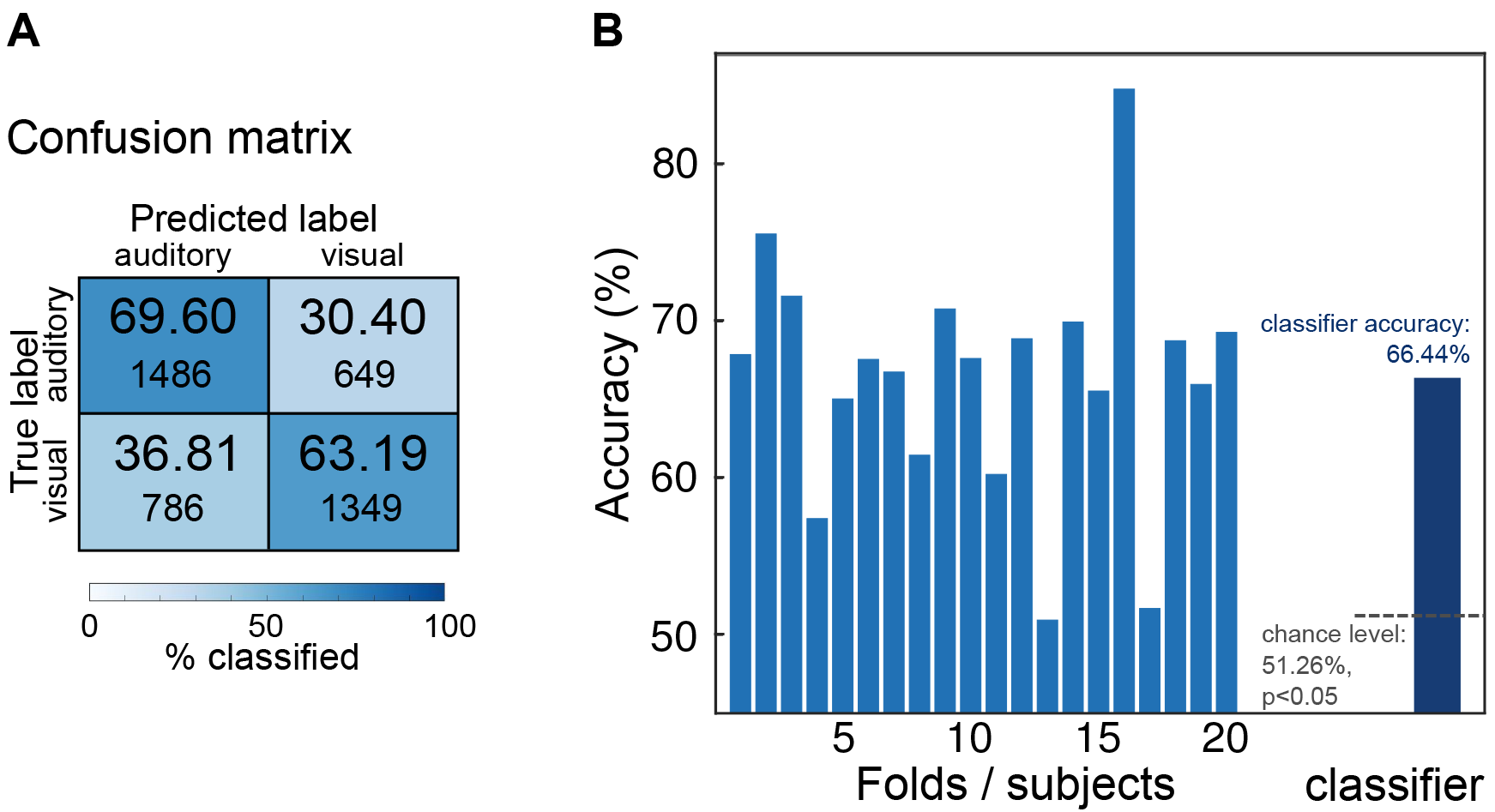
Classifier results. **A** Confusion matrix showing the proportion of correctly classified trials (diagonal) and misclassified trials. **B** Classifier accuracy. The mid blue bars represent the test accuracy in the 20 folds, where every fold is a subject the classifier was not trained on. The dark blue bar shows the overall classifier accuracy, which was tested against chance level.

The random forest classifier provides the variable importance as an importance estimate for every predictor in the model. This measure indicates the informational value of a given predictor towards the discrimination of the two classes, auditory and visual modality. Figure 2 reports the highest 2 % of variable importance values, i.e., those [voxel, time point, frequency band]-triplets that were most informative for partitioning the data. Not only visual, but also auditory regions contributed to the model, even though visual areas yielded more information than the auditory cortex. Interestingly, the lower frequency bands of 25-45 Hz and 55-75 Hz did not rank as important as the 75-95 Hz band. Even frequencies above 100 Hz contributed to the model in both visual and right auditory cortex. Gamma power beyond 125 Hz, however, did not add substantially to the classification model.

**Figure 2:**
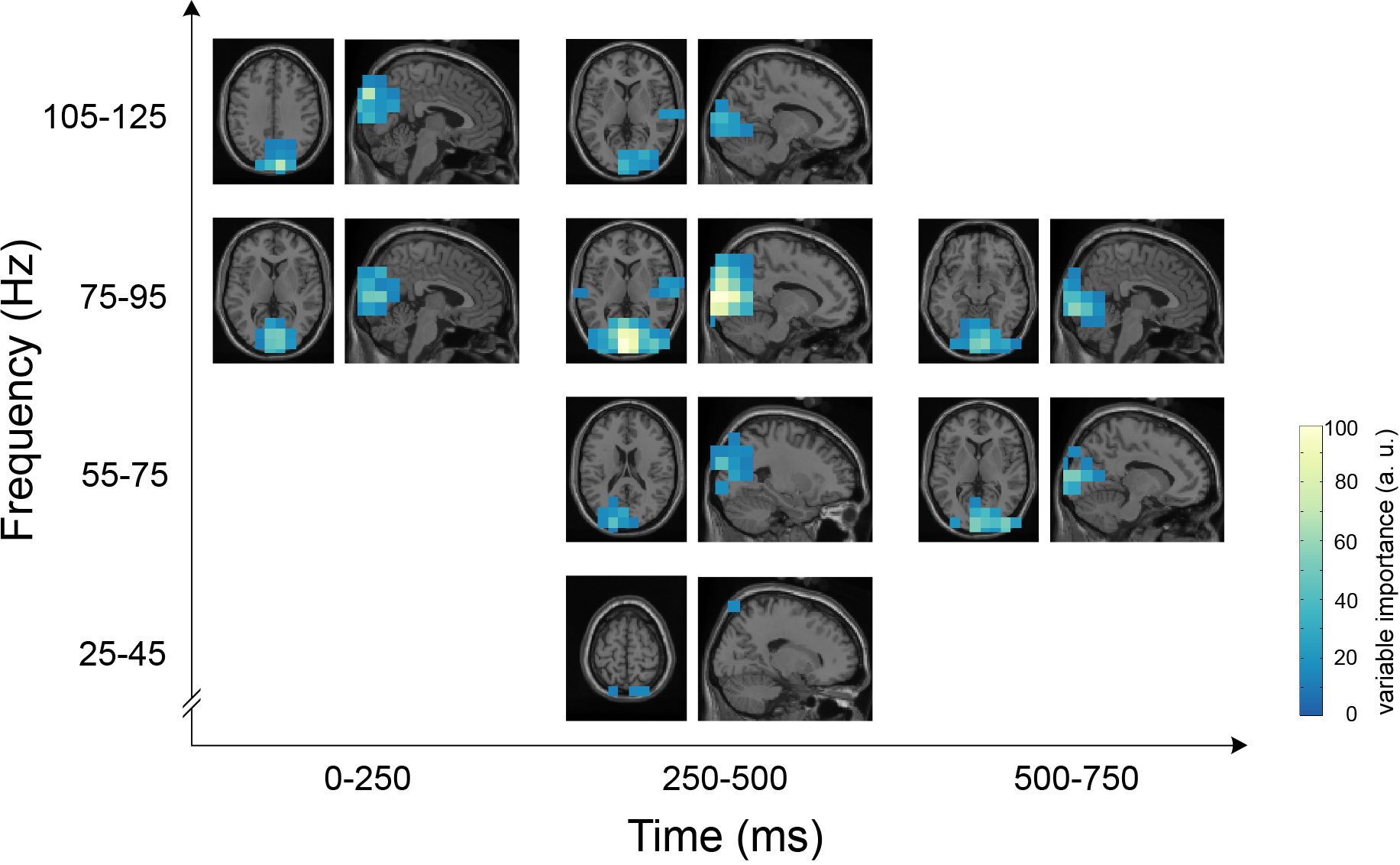
Variable importances. The 2 % most important predictors are shown across time, frequency and space. A higher variable importance score implies that this predictor had a higher informative value in the random forest model to partition the data into trials with auditory and visual perception. The orthogonal views are centered on the voxel showing the highest variable importance.

All time windows but the last one (750-1000 ms) supplied information to the classifier, in higher frequencies, the earlier time windows seemed to play a more pronounced role compared to the lower gamma frequencies. Figure 3 shows the time-frequency representations of variable importance for the visual and auditory peak voxels: the visual peak voxel (MNI coordinates: [−4 −100 12]) falls into left calcarine sulcus, the auditory peak voxel (MNI coordinates: [68 −20 10]) into right superior temporal gyrus (labels determined with the Automated Anatomical Labeling (AAL) atlas [25]). The time-frequency representations for those two peak voxels (Figure 3) confirm the pattern evident across all voxels (Figure 2). Thus, the 75-95 Hz band yielded a characteristic and stable activity pattern in both the auditory and visual cortex. The visual response was specifically characterized by a broadband gamma increase in the range of 55 to 125 Hz. The auditory response yielded informational value in an overlapping but narrower frequency range (75-125 Hz).

**Figure 3:**
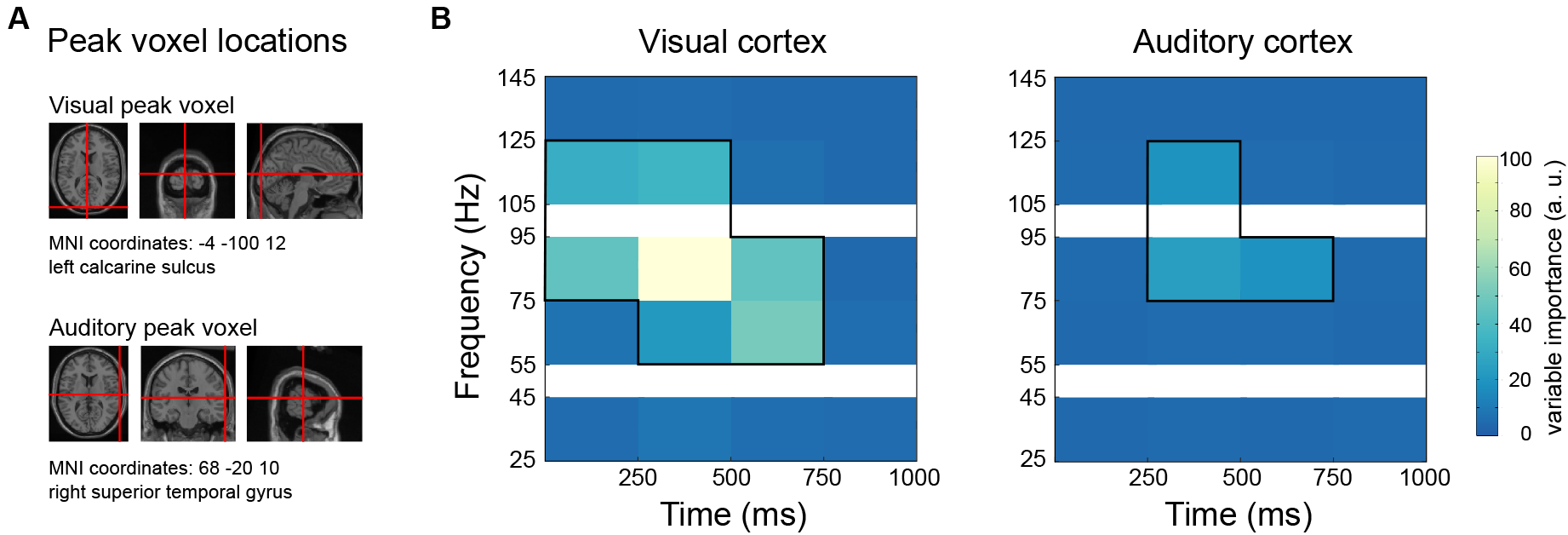
Variable importances in visual and auditory peak voxels. **A** Peak voxel locations for auditory and visual cortex (compare peak voxels from Figure 2) **B** Time-frequency representation of variable importances in those peak voxels. Black boxes indicate those variables which were among the 2% most informative predictors.

To investigate the underlying gamma power changes, the variable importance rankings were compared to the power differences between auditory and visual trials. To this end, auditory and visual power changes relative to baseline were averaged across trials and subjects, and the difference between the visual and the auditory condition was computed. These differences are depicted in Figure 4A: the spatial pattern of power is shown for the 250-500 ms time window and two frequency bands (75-95 Hz, top, and 105-125 Hz, bottom in Figure 4A). Red colors refer to higher gamma power in the visual condition and blue colors to higher power in the auditory condition. The black lines encircle those voxels which were among the 2% most informative predictors for the classifier. Figure 4B shows the underlying gamma power relations for the same peak voxels as presented in Figure 3. Interestingly, the classifier analysis based on single trials also rated predictors as highly informative where a difference in the averages is small, as is most evident for the time-frequency representation of the auditory condition (75-95 Hz, 500-750ms).

**Figure 4:**
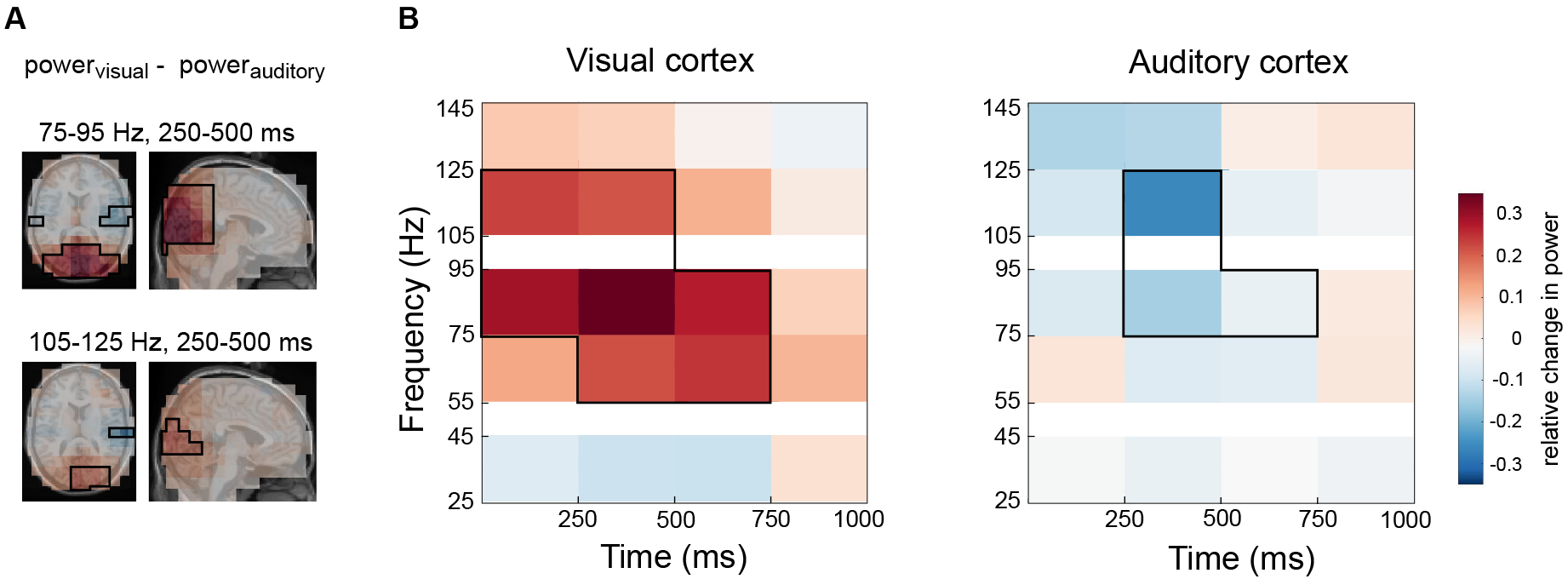
Underlying gamma power. This figure shows the difference in averaged gamma power between visual and auditory word presentation trials. **A** Spatial representation of gamma power for two frequency bands (75-95 Hz, top, and 105-125 Hz, bottom). Red hues represent a higher gamma power in the average of visual trials, the blue colors depict higher gamma power in the average of the auditory condition. Black boxes indicate the 2 % most informative predictors as shown in Figure 2. **B** Gamma power in visual and auditory peak voxels. Shown is the difference between the visual and auditory condition, black boxes again indicate the most informative predictors for the classifier model.

## Discussion

In the present work, we investigated the predictive value of single-trial gamma power to classify the stimuli’s modality. This was done in an across-subjects cross-validation framework which allowed us to estimate not only the gamma pattern stability across trials but also across subjects.

The decoding of MEEG high frequency activity on a single-trial basis can be challenging due to the low SNR: while intracranially recorded high frequency activity up to 180 Hz has been used to decode movements [26, 27], comparable approaches with MEEG data were not successful [28, 29]. Some studies could show a contribution of high gamma power (along with lower oscillatory activity) to the overall classifier performance [30, 31]. In this study, we successfully decoded stimulus modality exclusively from high frequency activity: the classifier model was able to correctly classify 66.44 % of the trials based on their source reconstructed gamma activity pattern, reliably distinguishing visual from auditory word presentation. Thus, the SNR of single-trial gamma power in source-level MEG data was high enough to successfully apply single-trial multivariate analyses. Interestingly, more auditory (69.30 %) than visual trials (63.19 %) were classified correctly, although visual areas yielded more information to the classifier. One possible explanation for this could be that the classifier-inherent cutoff values for gamma power in the visual voxels were rather conservative and therefore missed small gamma increases in visual cortex in visual trials, but still reliably detected the absence of visual activity in auditory trials.

The classification model was built across subjects, adopting a 20-fold across-subjects cross-validation, where the classifier was trained on 19 subjects and then tested on the data of the left-out 20^th^ subject. Hence, the trials of any given subject were classified by a model which was built on the data from different subjects. Using this approach, we assessed the common patterns across trials and subjects. The accuracy pattern across the different folds was higher than 60 % for all but three subjects. Low accuracies indicate either higher noise levels in these participants or activity patterns which deviate from the across-subjects consensus as uncovered by the random forest model. The overall classification accuracy of 66.44 % is comparable to previous reports of across-subjects MEG data classification (e.g., [32]).

The variable importance indicates which predictors were used by the model to yield the classification performance, by providing the common pattern across trials and subjects that differentiated between the two conditions. Clearly, gamma band activity from both visual and auditory areas was exploited by the model, although the visual cortex was more important than the auditory cortices, expressed by higher ranking variable importances. Overall, a broad range of frequencies and a time span of 750 ms included gamma band activation relevant to the random forest model.

In this study, we show the feasibility of applying the random forest algorithm [24] to single-trial source-localized time-frequency data. With its non-parametric, non-linear approach and its capability to handle high dimensional datasets with highly correlated predictors, this method is well suited for MEG data (also see [33, 34, 35, 36]) and can detect subtle differences concealed in the averaged data.

Another advantage of this method is the possibility to directly compare predictors
(e.g., frequency bands) to each other regarding their importance in the model: for example, we are able to state that the 75-95 Hz frequency band is the most important frequency band, and that the visual cortex has higher informational value for the classification than the auditory cortex.

In our data, the left primary visual cortex was most informative for the classification among all brain regions. Additionally, also higher visual areas ranked as highly informative, which is concordant with the localization of visual gamma band responses in intracranial electroencephalography (iEEG) and MEG studies (e.g., [37, 38, 5, 39]). The classification further identified auditory regions as informative. Although iEEG reliably shows high gamma responses to auditory stimuli [40, 41, 42, 43], auditory high frequency activity above 75 Hz has only rarely been shown in MEG studies: examples include high gamma responses to sound and pitch perception [44, 45]. Within the auditory regions, the most important voxel in our data was located in the right superior temporal gyrus, which is in line with iEEG studies investigating phoneme and word processing [7, 43] and the above-mentioned MEG studies. Further important regions included Heschl’s gyrus and the planum temporale. Interestingly, the right auditory cortex showed higher importance with more voxels involved compared to the left auditory cortex, although the stimuli were words and should typically evoke language-related activity localized to the left hemisphere [42, 43]. This may be explained by the fact that both conditions used words as stimuli and thus, left-hemispheric language related activity is not able to distinguish between auditory and visual trials.

The time windows most important to the classification covered 0 to 750 ms after stimulus onset, while the last time window (750-1000 ms) did not show any high ranking variable importance values, implying that gamma activity was most informative to the classifier during presentation of a word (mean = 700 ms).

Informative predictors in auditory areas, however, were only found between 250 and 750 ms, although previous studies reported early auditory (high) gamma responses following phoneme or word stimuli (e.g., [7, 43, 46, 47]).

In both, visual and auditory brain areas, the most important frequency band was the 75-95 Hz band. Especially in the 250-500 ms window, this frequency band exhibited exceeding informative value for the classification. Yet, the visual areas overall provided informative predictors across a broad frequency range (55-125 Hz). This points to underlying broadband gamma activity in single trials rather than a narrowband response, which is typically elicited by high contrast stimuli such as gratings, (e.g., [5, 39]). The high frequency activity beneficial for classification is similar to visually induced broadband gamma activity reported in iEEG and MEG studies [48, 37, 49, 50, 6]. Vidal et al. [49], for example, describe a lower frequency band of 45-65 Hz and high gamma activity of 70-120 Hz in their MEG study on visual grouping. Related to reading, broadband high frequency activity above 50 Hz has been reported in iEEG studies [51, 52, 53, 54]. Furthermore, compared to the narrowband responses elicited by high contrast stimuli such as gratings, which are typically centered at lower frequencies, (e.g., 50 Hz [5] or 60 Hz [39]), our results yielded the 75-95 Hz frequency band as most informative.

In the auditory areas, the most important variables were concentrated in the 75– 95 Hz and 105-125 Hz Hz frequency bands. This is in line with iEEG studies on syllable and word processing, which report gamma responses from 80 Hz up to 200 Hz [42, 55]. Thus, our results might reflect the lower end of the high gamma response described in these studies, potentially cropped by low SNR above 125 Hz.

To summarize, we have shown that single-trial gamma activity can be successfully used to classify stimulus modality. Importantly, the successful across-subjects classification suggested that single-trial gamma-band activity contains high inter-individual consistency. The classifier identified both visual and auditory areas as informative with high spatial specificity. Our results furthermore suggest that single-trial high frequency activity after visual word presentation is characterized by a broadband rather than a narrowband response.

## Methods

### Ethics statement

The study was approved by the Institutional Review Board of the University of Konstanz and in accordance with the Declaration of Helsinki.

### Participants

A total of 24 participants (17 female; mean age=22 years, range= 19-26 years; 21 right-handed) took part in this MEG experiment. Three participants were excluded due to technical problems, one due to excessive environmental noise. The data from the remaining 20 participants are presented here. All of the participants gave written informed consent prior to the experiment and received course credits or nominal financial compensation for participation. All participants were German native speakers and reported normal or corrected-to-normal vision, and no history of neurological disease.

Parts of this data have been published in [12], with respect to independent research questions and analyses.

### Design, procedure, and material

The experiment consisted of a study phase and a subsequent recognition test. Only data from the study phase are reported here. In the study phase, participants were presented with words either visually (projected centrally on a screen) or auditorily (via nonferromagnetic tubes to both ears). The duration of the visual word presentation was determined by the duration of the respective audio file, i.e., the time to pronounce the word (mean duration = 697 ms, *s.d.* = 119 ms). Each word was followed by a fixation cross. The duration of the word and fixation cross together added up to 2000 ms. Participants were instructed to count the syllables of the word and indicate via button press whether the word had two syllables. A question mark (max. duration of 1500 ms) prompted the subject’s response. The button press ended the presentation of the question mark. A fixation cross with variable duration (1000-1500 ms) was presented before each item. After the encoding phase, participants performed a distractor task and a surprise recognition test phase.

The stimuli consisted of 420 unrelated German nouns, grouped into three lists with 140 words. Half of each list’s words had two syllables, the other half had one, three or four syllables. Two lists were presented during the study phase and one list during the test phase. The assignment of the lists to study or test phase was counterbalanced across participants. Items were presented in random order, with the constraint that not more than 5 words of the same modality and not more than 5 words from the same condition were presented sequentially.

### MEG data acquisition and preprocessing

MEG data was recorded with a 148channel magnetometer (MAGNES™ 2500 WH, 4D Neuroimaging, San Diego, USA) in a supine position inside a magnetically shielded room. Data was continuously recorded at a sampling rate of 678.17 Hz and bandwidth of 0.1-200 Hz, and later downsampled to 300Hz to reduce computational load. All data processing prior to classification was done using FieldTrip [56], an open-source MATLAB toolbox for MEEG data analysis. Data was epoched into single trials, with epochs ranging from 1500 ms before item presentation to 4000 ms after item presentation. Trials were visually inspected for artifacts, contaminated trials were rejected. Thereafter, trials were corrected for blinks, eye movements, and cardiac artifacts using independent component analysis (ICA).

### Source reconstruction

For coregistration with the individual structural magnetic resonance image (available for 17 out of 20 participants; for the remaining three participants we used an affine transformation of an MNI-template brain; Montreal Neurological Institute, Montreal, Canada), the shape of the participant’s head as well as three markers (nasion, left and right ear canal) and the location of the head position indicator (HPI) coils were digitized prior to the experiment using a Fastrak Polhemus 3D scanner (Polhemus, Colchester, VT, USA).

Single-trial source space activity was reconstructed using a LCMV beamformer [23] with weight normalization (neural activity index; [23, 57]). First, the spatial filter was computed adopting a realistic single shell head model [58] based on the individual structural magnetic resonance image (MRI) and a source model with grid points covering the whole brain volume (resolution: 15 mm). The data covariance matrix was computed for −500 to 1000 ms relative to stimulus presentation. Subsequently, the spatial filter was applied to the single trials to obtain virtual electrodes at all grid point locations.

For the classification of the oscillatory activity, single-trial time frequency representations were calculated at every virtual electrode applying a Fast Fourier Transform. Gamma band activity was estimated using frequency smoothing (Slepian sequence multi taper approach), yielding 20 Hz-wide frequency bands centered at 35, 65, 85, 115 and 135 Hz. The power was calculated separately for 250 ms long time windows from −500 to 1000 ms and the post-stimulus activity was then expressed as relative change to baseline power.

### Random forest classification

The random forest algorithm [24], an ensemble method, aggregates the results of several classifiers. These so-called base learners are classification and regression trees [59], which partition the data by adopting binary splits. The aim of this partitioning process is to reduce the impurity regarding the class labels in the daughter nodes that result from this split: preferably, all observations from one class should arrive in the same node. In every split, the tree algorithm searches first for the predictor that maximizes the purity of the daughter nodes and then for the best split point within that predictor. Random forests now grow numerous trees; each of these trees, however, is built on a bootstrap sample of the original data and in every split only a random subsample of all predictors is searched. The variance introduced by this randomness leads to a robust prediction by the aggregated model. This approach furthermore enables random forest to cope particularly well with highly correlated predictor variables [60], which is of special interest when working with MEEG data.

Additionally, data with more predictors than observations (small *n* large *p* problems) are also handled effectively since the predictor variables are searched successively [61], which makes this approach particularly interesting when dealing with high-dimensional source-space MEEG data. For every predictor, the algorithm returns an estimate of how important this variable was for the model’s prediction. The version used here is based on the impurity reduction introduced by a predictor variable across all trees, which is measured by the Gini index [62, 59, 60].

Random forest classification was performed using the scikit-learn module for Python [63]. The aim of the decoding was to classify trials regarding their stimulus modality: visual or auditory. The predictors were [voxel, time point, frequency band]-triplets, providing 16 624 predictors, overall. For every subject, the more prevalent class (auditory or visual stimulation) was downsampled such that every dataset contained equal trial numbers for both cases. The total trial number across all subjects was 4270 trials.

The classification was embedded in a cross-validation framework across subjects: the classifier was trained on the data from all but one subject and then tested on the data of this left-out subject. This procedure was repeated for all 20 subjects, such that every dataset was used as test set once. This approach ensures that the classifier is never tested on data it was trained on and thus controls for possible overfitting of the classifier. Moreover, it allows the assessment of across-subjects predictability of the data regarding the response variable.

Each of the 20 cross-validation models aggregated the results of 15 000 classification trees, where every tree was built on a bootstrap sample of all observations in the trainings set. To ensure that the model incorporated a sufficient number of trees, classification performance was assessed with 25 000 trees for two folds [64], yielding comparable results as the sparser model. At each binary split, the algorithm considered 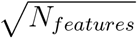 predictors to find the best split. The accuracies on the test datasets as well as the variable importances were merged across the cross-validation folds. The performance of the classifier was then tested against 50% chance level using a binomial test [65], since a permutation based test was computationally not feasible.

## Acknowledgements

**Acknowledgments** We thank Ann-Kristin Rombach, Leona Hellwig, Marina Koepfer, and Janine Weichert for help with data acquisition and Sabine Leske and Tzvetan Popov for valuable discussion. This project was supported by ERA-Net NEURON via the German Federal Ministry of Education and Research (BMBF, grant 01EW1307 to SSD) and the European Research Council (Starting Grant 640488 to SSD). This research was further supported by an Emmy Noether Programme Grant from the Deutsche Forschungsgemeinschaft awarded to SH (HA 5622/1-1). SH is supported by a European Research Council Consolidator Grant (Grant Agreement 647954), the Wolfson Society and the Royal Society. TS is supported by funding from the European Union’s Horizon 2020 research and innovation programme under grant agreement No 661373.

